# Electrostatic Interactions Dictate Bile Salt Hydrolase Substrate Preference

**DOI:** 10.1101/2023.09.25.559308

**Authors:** Kien P. Malarney, Pamela V. Chang

## Abstract

The human intestines are colonized by trillions of microbes, comprising the gut microbiota, which produce diverse small molecule metabolites and modify host metabolites, such as bile acids, that regulate host physiology. Biosynthesized in the liver, bile acids are conjugated with glycine or taurine and secreted into the intestines, where gut microbial bile salt hydrolases (BSHs) deconjugate the amino acid to produce unconjugated bile acids that serve as precursors for secondary bile acid metabolites. Among these include a recently discovered class of microbially-conjugated bile acids (MCBAs), wherein alternative amino acids are conjugated onto bile acids. To elucidate the metabolic potential of MCBAs, we performed detailed kinetic studies to investigate the preference of BSHs for host- and microbially-conjugated bile acids. We identified a BSH that exhibits positive cooperativity uniquely for MCBAs containing an aromatic sidechain. Further molecular modeling and phylogenetic analyses indicated that BSH preference for aromatic MCBAs is due to a substrate-specific cation-ρε interaction and is predicted to be widespread among human gut microbial BSHs.

**TOC graphic:** 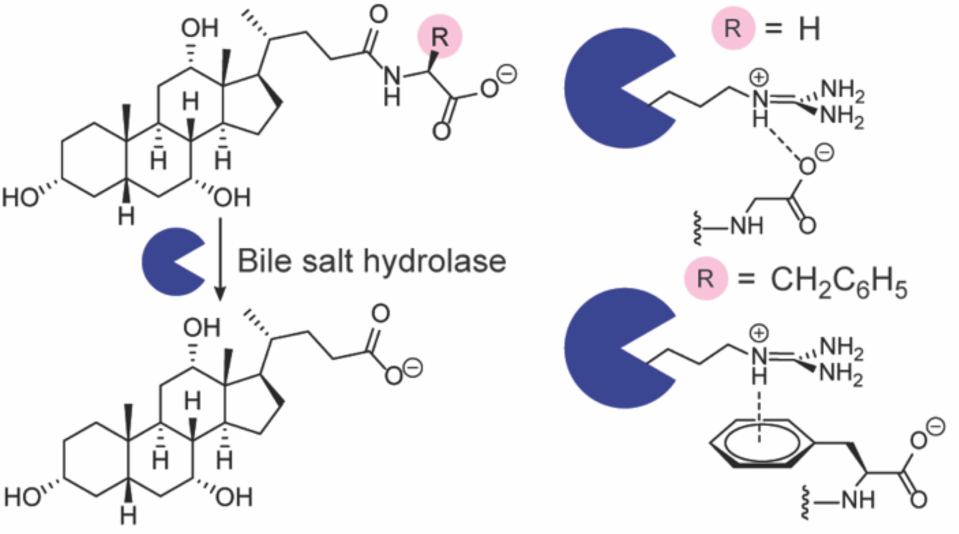

## Introduction

Humans are colonized by a dynamic and diverse community of 100 trillion microorganisms termed the human microbiome. Microbes residing in the intestines, known as the gut microbiota, carry out important metabolic functions, including breakdown of otherwise indigestible dietary compounds and production of nutrients and signaling molecules that regulate host physiology.^1^ Bile acids (BAs) are a key class of metabolites whose function is to aid in the absorption of fat-soluble, dietary vitamins and lipids and undergo extensive metabolism by the gut microbiota.

These molecules also serve as signaling agents that activate host receptors and are important regulators of host metabolism^2^ and immunity^3^. In addition, BAs function as antimicrobial agents against susceptible microbes and therefore impact gut microbial composition, which is correlated with host health.

BAs are synthesized from cholesterol in the liver, where they undergo conjugation with the amino acids glycine or taurine to increase water solubility (Fig. 1a). These host-conjugated BAs are postprandially secreted as emulsifying agents into the small intestine, where they encounter gut microbial bile salt hydrolases (BSHs) that catalyze amino acid deconjugation to produce unconjugated BAs (Fig. 1a).^4^ Gut microbial enzymes can then further metabolize unconjugated BAs through epimerization and redox modifications to produce secondary BAs.^4^ Recent work demonstrated that unconjugated BAs can also be reconjugated to a wide variety of many amino acids, including tyrosine, phenylalanine, and leucine, by microbial enzymes to produce microbially-conjugated bile acids (MCBAs, Fig. 1a).^5–11^ As BSH activity is thought to precede all subsequent BA transformations, this enzyme has been termed the “gatekeeper” enzyme of secondary BA metabolism.^2,7^

**Figure 1.**
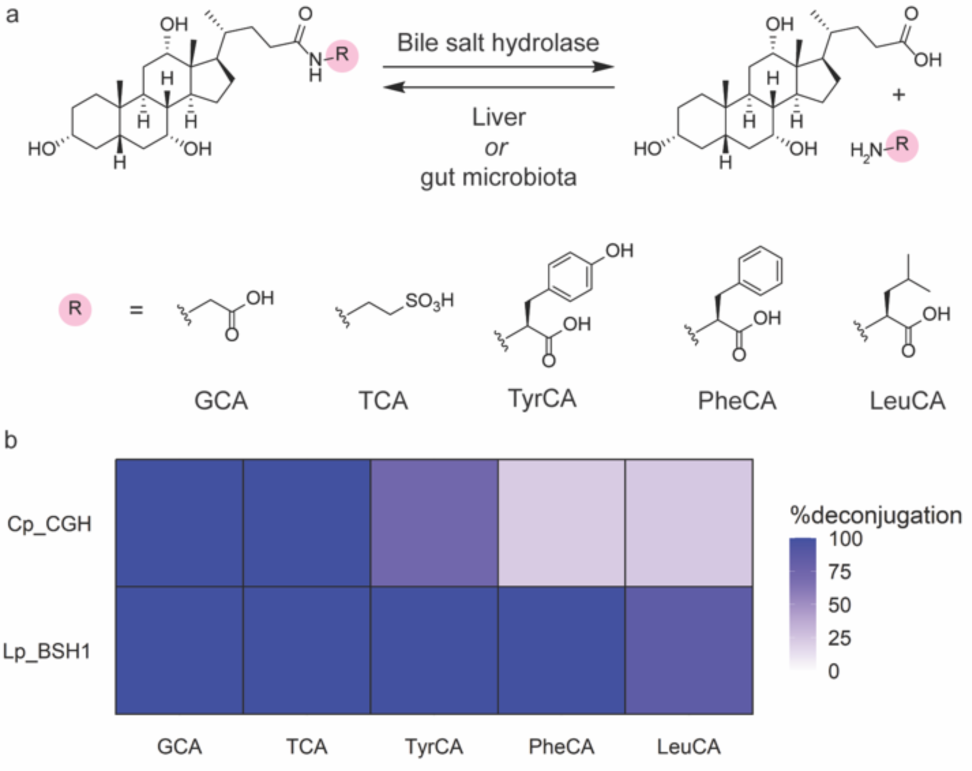
(a) Gut microbial bile salt hydrolases (BSHs) catalyze the hydrolysis of bile acid conjugates, whereas amino acid conjugation is performed by liver or gut microbial enzymes. (b) Activity heat map of host- and microbially-conjugated bile acid hydrolysis by *C. perfringens* CGH (Cp_CGH) and *L. plantarum* BSH1 (Lp_BSH1). BSH (387.8 nM) was incubated with 1 mM bile acid in phosphate buffered saline, pH = 6.2, with 10 mM DTT for 16 h at 37 °C. Data are representative of three independent experiments.

Previous work has shown that BSHs have distinct substrate specificities, determined by both the identity of the conjugated amino acid^12,13^ and the steroid core of the bile acid.^2,14,15^ A recent report by Theriot and co-workers demonstrated that BSH side chain preference for the host-conjugated BAs is determined by a selectivity loop containing a three amino acid motif.^16^ They further showed that the newly discovered MCBAs also serve as substrates for BSHs and alter host BA pools that impact the host during infection.^16^

Given the importance of BAs in human health, we sought to biochemically investigate the BSH substrate preferences for MCBAs to understand how its activity in deconjugating these novel BAs contributes to the host BA pool. Here, we utilize biochemical in vitro assays using purified BSHs from anaerobes present in the human gut microbiome, *Clostridium perfringens* and *Lactiplantibacillus* (formerly *Lactobacillus*) *plantarum*, to determine kinetic parameters and substrate specificities of several MCBAs containing side chains with unique chemical properties. We found that both BSHs preferred smaller, less sterically hindered amino acid side chains, whereas *L. plantarum* BSH1 exhibited some preference for MCBAs over host-conjugated BAs and positive cooperativity for aromatic MCBAs. From molecular modeling studies, we found that the previously characterized three amino acid selectivity loop^16^ also dictates BSH1 substrate preference of aromatic MCBAs through a larger four amino acid motif and a cation-ρε interaction. We find that this selectivity loop is widespread in many genera throughout the human gut microbiome, suggesting that BSH MCBA deconjugation is a widespread activity that contributes to secondary BA metabolism in the intestines.

## Materials and Methods

### General materials and methods

Chemicals were used as received unless otherwise noted. ACS-grade acetone and ethyl acetate (EtOAc) was obtained from Fisher Scientific (Waltham, MA, USA). Cholic acid was obtained from Beantown Chemicals (Hudson, NH, USA). Sodium taurocholate hydrate was obtained from Alfa Aesar (Haverhill, MA, USA). Sodium glycocholate hydrate and ethyl chloroformate were obtained from Sigma-Aldrich (St. Louis, MO, USA). Triethylamine was obtained from Chem-Impex (Wood Dale, IL, USA). L-leucine, L-phenylalanine, and L-tyrosine were obtained from Fluka Chemical Corporation (Buchs, Switzerland). Flash chromatography was performed using Siliaflash P60 40-63Å 230-400 mesh silica gel obtained from SiliCycle (Quebec City, Canada). Thin layer chromatography (TLC) was performed using glass-backed TLC 60 Å silica gel plates (Merck), with visualization by UV light (254 nm), followed by staining with cerium ammonium molybdate. Millex-LH syringe filters (0.45 μm PTFE membrane) were obtained from Millipore (Burlington, MA, USA). HisPur Ni-NTA resin (50% slurry) was obtained from Thermo Scientific (Waltham, MA, USA). LC-MS grade water, formic acid, and methanol were obtained from Fisher Scientific (Waltham, MA, USA). LC-MS grade acetonitrile was obtained from Millipore (Burlington, MA, USA).

### Instrumental analysis

NMR spectra were acquired on a Bruker-500 spectrometer. LC-MS analysis was performed using an Agilent 6320 electrospray ionization time-of-flight MS coupled to an Agilent 1260 HPLC fitted with an Agilent InfinityLab Poroshell 120 EC-C18 column (3.0 x 50 mm, 2.7 μm mesh) with a flow-rate of 0.5 mL/min. Bile acids were separated using a binary gradient elution system of water containing 0.1% formic acid (solvent A) and acetonitrile containing 0.1% formic acid (solvent B). Briefly, the column was eluted with 5% solvent B for 0.2 min, after which solvent B was increased linearly over 5.4 min to 95% solvent B. Solvent B was then linearly increased to 100% solvent B, followed by isocratic elution for an additional 2 min. Bile acids were detected using an Agilent Jet Stream ionization source, acquiring in extended dynamic range from m/z 110-1050 in negative mode at 3 spectra/s; fragmentor voltage: 300 V; nozzle voltage: 1000 V; Vcap: 4000 V; nebulizer: 35 psig; gas temperature: 200 °C; drying gas: 12 L/min; sheath gas flow: 12 L/min; and sheath gas temperature: 300 °C.

### Kinetic and statistical analyses

Regression analyses were performed in RStudio (version 2022.12.0.353). Reported values are representative of three independent experiments. Error is reported as the standard error of the fitted value or appropriate propagated error.^17^ Saturating kinetics were fitted to the Michaelis-Menten equation or Hill equation^18^ using nonlinear least squares, for which initial parameters were estimated from visual inspection of kinetic plots. In both cases, *kcat* was determined as the ratio of *Vmax* and the enzyme concentration. Non-saturating kinetics were analyzed by linear regression. Briefly, *Vmax* values were calculated from the maximum fitted value, and *kcat* was determined as the ratio of *Vmax* and the enzyme concentration. *Km* values were calculated as the fitted concentration at which the observed half-maximal velocity was obtained. Differences in mutant and wildtype enzyme activity were compared using Welch’s t-test (paired).

### Molecular modeling analysis

The structure of *L. plantarum* BSH1 was generated using ColabFold-AlphaFold2.^19^ Briefly, the protein was sequence (NCBI accession: CCC80500.1) was truncated to have a N-terminal cysteine^20^ and inputted as a homotetramer.^21^ Structure prediction parameters were left unaltered.

Docking analyses were performed in Maestro (version 13.1.137; release 2022-1; Schrödinger, New York, NY, USA). Protein and ligand preparation was performed using ProteinPrep and LigPrep, respectively.^22^ The protonation state of the *L. plantarum* BSH1 Cys2 thiol was manually assigned to a thiolate.^20^ Receptor grids were generated with a 12 Å inner box and 28 Å outer box, using a 5 Å distance constraint for the Cys2 sulfur^20^ atom. The center of each grid was placed at the center of each monomer active site, with coordinates [33, 5, 24] and [-23, 17, 2] for *C. perfringens* CGH and *L. plantarum* BSH1, respectively. Ligand docking was then performed with Extra Precision Glide,^23^ with the constraint that 1 atom of the bile acid conjugate amide must be within 5 Å of the Cys2 thiolate. The docked pose of PheCA was used to generate template-based models for LeuCA and TyrCA complexes with *L. plantarum* BSH. In Maestro, the appropriate phenyl ring atoms were deleted and modified to an isopropyl group for LeuCA, and a *para*-hydroxyl group was added for TyrCA. For *C. perfringens*, appropriate modifications were used to generate template-based models for PheCA and LeuCA using the obtained docking pose for TyrCA. Models obtained from docking were then minimized using Prime^24^ and visualized in VMD.^25^ Cation-π interactions were identified using previously described geometric analysis.^26^

### Bioinformatic analysis

Protein sequences of the cholylglycine hydrolase subfamily (EC 3.5.1.24) were obtained from the NCBI Protein database with a length cutoff of 400 amino acids.^27^ Taxonomic assignments were then obtained from the NCBI Taxonomy database^28^ using taxize.^29^ Taxa were then analyzed in the NCBI BioSample database for their presence in the human gut, and only sequences from human gut-derived taxa were used for further analysis. Protein sequences were analyzed for the presence of “X1-S-R-X2,” where X1 = Y, F, or W and X2 = G, S,^16^ or A. Due to the presence of leader sequences,^30^ sequences were filtered according to the presence of this motif beginning between the 200^th^ and 230^th^ position. For phylogenetic analysis, protein sequences were aligned using Clustal Omega.^31^ Phylogenetic tree generation was performed using BLOSUM62^32^ for distance scoring and neighbor-joining in phangorn^33^ and visualized with FigTree. For genus-level analysis, sequences without genus-level assignments were omitted from the analysis.

### Plasmids and cloning

A previously described protein expression plasmid for CGH from *Clostridium perfringens* 13 (NCBI accession: P54965.3) was used in this study.^34^ BSH1 from *Lactiplantibacillus plantarum* WCSF1 was cloned using an analogous approach. Briefly, *bsh1* from *L. plantarum* WCSF1 was PCR-amplified from a synthetic gene fragment (gBlocks, Integrated DNA Technologies, San Diego, CA, USA). Amplification using a forward primer that introduced a NdeI restriction digest site (CTAGCATATGTGTACTGCTATCACCTATCAGTCTT) and reverse primer that introduced a C-terminal FLAG tag (DYKDDDDK) and XhoI restriction digest site (CTAGCTCGAGCTTATCGTCGTCATCCTTGTAATCGTTAACTGCATAGTATTGTGCTTCT) generated an amplicon that was cloned into a pET21b plasmid containing an ampicillin selection marker.

Site-directed mutagenesis was performed using the QuikChange kit (Agilent Technologies, Santa Clara, CA, USA). For *L. plantarum* BSH1, the R207Q mutant was generated using a mutagenic forward primer (GTGGATTTAGATAGTTATAGTCAAGGAATGGGCGGACTAGG) and reverse primer (CCAGGTAATCCTAGTCCGCCCATTCCTTGACTATAACTATC). For *C. perfringens* CGH, the Q212R mutant was generated using a mutagenic forward primer (GACGGCTCTTGGACGGGGCACGGGTTTAG) and reverse primer (CTAAACCCGTGCCCCGTCCAAGAGCCGTC).

### Protein expression and purification

Protein expression and purification of BSHs was performed according to literature procedure.^30,34,15^ Briefly, a freshly-transformed plate of *E. coli* Rosetta 2 (DE3) pLysS competent cells on L-agar containing ampicillin was used to inoculate L-broth containing chloramphenicol and ampicillin, which was cultured with shaking at 37 °C until an OD600 of 0.5-0.6 was reached. The culture was then transferred to Terrific Broth (12 g/L Tryptone, 23.9 g/L yeast extract, 8 mL/L glycerol, 0.22 g/L KH2PO4, 0.94 g/L K2HPO4) containing chloramphenicol and ampicillin, which was subsequently grown with shaking at 37 °C until OD600 of 0.5-0.6 was reached. Protein expression was then induced with 0.1 mM IPTG with shaking at 18 °C for 18 h.

The culture was then aliquoted to 50 mL conical tubes on ice, and cells were pelleted by centrifugation (4696 x *g*, 4 °C, 30 min). The supernatant was removed, and the cells were resuspended in lysis buffer (PBS (pH 7.4) + 5% glycerol containing 250 μM TCEP, 1 mM PMSF, and 20 mM imidazole). The cells were pelleted once more (4696 x *g*, 4 °C, 10 min), and the cells were resuspended in lysis buffer. Lysis was then performed by sonication at 0 °C, and the soluble fraction of the lysate was then isolated by centrifugation (4696 x *g*, 4 °C, 30 min).

During this time, Ni-NTA beads were equilibrated by washing with PBS (pH 7.4) + 5% glycerol containing 250 μM TCEP, and 20 mM imidazole (3x). The bacterial lysate was then incubated with the Ni-NTA beads for 90 min at 4 °C with nutation. The flow-through was then drained, and the beads were washed with PBS (pH 7.4) + 5% glycerol containing 250 μM TCEP, and 20 mM imidazole (20x). The protein was then eluted with PBS (pH 7.4) + 5% glycerol containing 250 μM TCEP, 1 mM PMSF, and 500 mM imidazole (4x). The elutions were then concentrated on an Amicon spin column (10 kDa cutoff) and quantified by the DC assay (Bio-Rad, Hercules, CA, USA). The eluted protein was then diluted in PBS (pH 7.4), aliquoted, flash-frozen in liquid nitrogen, and stored at -80 °C until further use. Purity was assessed by SDS-PAGE.

### Kinetic studies

An aliquot of BSH was thawed on ice and diluted in PBS (pH 6.2). Assays were performed in PBS (pH 6.2) containing 12.5 mM DTT (assay buffer). For time-course experiments, 88 μL of assay buffer and 11 μL of conjugated bile acid (BA, 10x stock in DMSO) were added to a 1.7 mL microfuge tube and mixed. An aliquot (9 μL) was then quenched with 6 M HCl in a 1.7 mL microfuge tube as an initial timepoint. Otherwise, 80 μL of assay buffer and 10 μL of conjugated bile acid (10x stock in DMSO) were added to a 1.7 mL microfuge tube and mixed.

The BSH reaction solutions and enzyme solutions were then prewarmed to 37 °C, and 10 μL of BSH stock was added to create a 100 μL reaction (10 mM DTT, 1x BA). At each time point, a 10 μL aliquot was removed and quenched with 90 μL of 6 M HCl in a 1.7 mL microfuge tube. The aqueous layer was then extracted with EtOAc (140 μL), and the phases were separated by centrifugation (17,900 x *g*, 1 min). The organic layer (100 μL) was transferred, and the aqueous layer was extracted with EtOAc once more. The combined organic extracts were then dried under a stream of nitrogen, and the resulting residue was dissolved in LC-MS grade MeOH (1 mL). The resulting solution was then taken up in a gas-tight syringe, filtered through a 0.45 μm PTFE membrane, and subjected to LC-MS analysis with three technical replicates. Quantification of bile acids was determined by integrating the extracted ion count of the exact masses of the bile acids compared to standard curves of synthetic and commercial standards. Percent deconjugation was calculated as the ratio of unconjugated bile acids and total bile acids^2,30^ detected after correcting for background hydrolysis from a PBS blank reaction. For comparison of substrate conversion between enzymes, single timepoint reactions were used for bile acid quantitation after correcting for background hydrolysis from a PBS blank reaction.

### Synthetic procedures

TyrCA, PheCA, and LeuCA were synthesized according to literature procedures.^5^

## Results and Discussion

To test the hypothesis that BSHs deconjugate MCBAs, we utilized cholic acid-derived MCBAs that have been identified in human fecal samples, which are conjugated to tyrosine, phenylalanine, or leucine (TyrCA, PheCA, LeuCA; Fig. 1a).^5^ Due to the importance of the steroid core in BSH substrate preference,^2,15^ we characterized the activity of BSHs that are known to hydrolyze CA conjugates, including CGH^35^ from *C. perfringens* 13 and BSH1^14^ from *L. plantarum* WCSF1. *L. plantarum* BSH1 was recombinantly expressed as previously described for *C. perfringens* CGH (Fig. S1).^34^ We then characterized their ability to hydrolyze MCBAs, along with host-derived BA conjugates, glycocholate (GCA) and taurocholate (TCA), by LC-MS analysis. Indeed, incubation of TyrCA, PheCA, and LeuCA with these BSHs for 16 h revealed modest deconjugation of MCBAs by *C. perfringens* CGH, as well as extensive deconjugation by *L. plantarum* BSH1 (Fig. 1b).

Although previous literature suggested *C. perfringens* CGH hydrolyzes LeuCA more readily than TyrCA,^13^ we found TyrCA was hydrolyzed to a greater extent than LeuCA (Fig. 1b) because *C. perfringens* show strain-dependent substrate preferences based on the identity of the amino acid conjugate.^36,37^ We also found that TyrCA and PheCA had remarkably different amounts of hydrolysis despite comparable steric hindrance near the amide bond, which was a previously identified factor for substrate preference^13^ (Fig. 1b). These differences in overall MCBA hydrolysis between CGH and BSH1 suggested distinct substrate preferences (Fig. 1b). To understand these results, we characterized BSH substrate preference using kinetic studies.

We first determined conditions for which hydrolysis was linear over time for CGH (Fig. S2) and BSH1 (Fig. S3). Kinetic characterization revealed distinct substrate preferences between CGH and BSH1 (Fig. 2). CGH exhibited classical Michaelis-Menten kinetics for host-conjugated BAs (Fig. 2a-b) with TCA having a higher turnover number, lower *Km* value, and a higher catalytic efficiency than GCA (Table 1). The *Km* values fell within the range of known values for CGH expressed by different strains of *C. perfringens*.^37^ Previous characterization of *Escherichia coli* lysates overexpressing CGH also identified this BSH as preferring taurine-conjugated BAs.^35^ MCBAs were recalcitrant to hydrolysis by CGH and exhibited linear, non-plateauing kinetics (Fig. 2c-e). As such, kinetic parameters of CGH for MCBAs were determined by linear regression and were substantially lower than those for TCA and GCA (Table 1).

**Figure 2.**
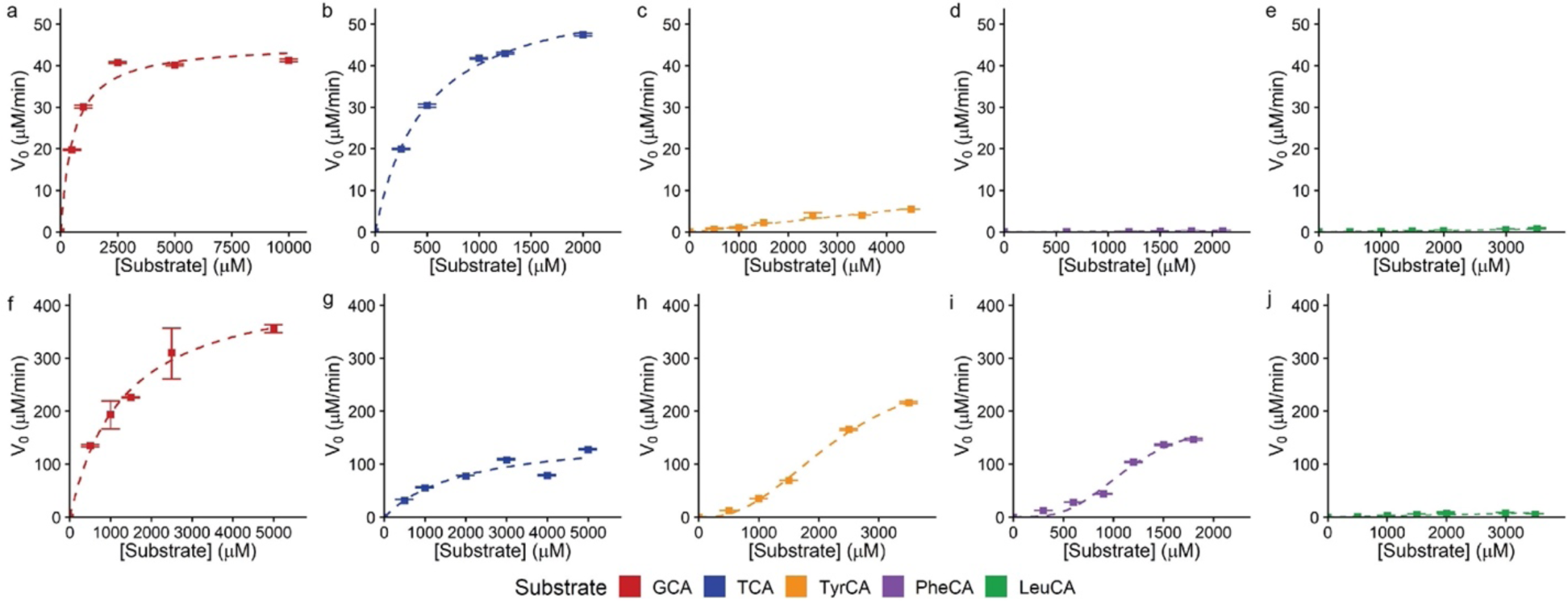
Kinetic characterization of *C. perfringens* CGH (a-e) and *L. plantarum* BSH1 (f-j). BSH (a-b) 2.7 nM, (c-e) 387.8 nM, (f) 9.7 nM and (g-j) 193.4 nM was incubated with the indicated amount of substrate in phosphate buffered saline, pH = 6.2, with 10 mM DTT under initial rate conditions (a,b: 5 min; c-e: 1 h; f-i: 3 min; j: 10 min) at 37 °C. Data points were fit to the Michaelis-Menten equation (a-b, f-g), Hill equation (h, i), or linear regression (c-e, j). Data are representative of three independent experiments, n=3, points = mean, error bars = standard deviation.

**Table 1.** Kinetic parameters for BSHs from *C. perfringens* and *L. plantarum*.

| Enzyme | Substrate | $k_{cat}$<br>(min<sup>-1</sup>) | $K_m$<br>(μM) | $k_{cat}/K_m$<br>(min<sup>-1</sup>μM<sup>-1</sup>) | Hill coefficient<br>(N) |
| --- | --- | --- | --- | --- | --- |
| *C. perfringens*<br>CGH | GCA<sup>a</sup> | 16789 ± 392 | 540 ± 57 | 31.1 ± 3.4 | - |
|  | TCA<sup>a</sup> | 22074 ± 317 | 477 ± 20 | 46.3 ± 2.1 | - |
|  | TyrCA<sup>b</sup> | 14.7 ± 0.5 | 2169 ± 70 | 0.0068<br>± 0.0003 | - |
|  | PheCA<sup>b</sup> | 0.62 ± 0.03 | 1007 ± 354 | 0.0006<br>± 0.0002 | - |
|  | LeuCA<sup>b</sup> | 1.70 ± 0.07 | 1937 ± 449 | 0.0009<br>± 0.0002 | - |
| *L. plantarum*<br>BSH1 | GCA<sup>a</sup> | 46474 ± 2727 | 1300 ± 198 | 35.8 ± 5.8 | - |
|  | TCA<sup>a</sup> | 804 ± 101 | 1918 ± 598 | 0.42 ± 0.14 | - |
|  | TyrCA<sup>c</sup> | 1547 ± 94 | 2347 ± 140 | 0.66 ± 0.07 | 2.5 ± 0.1 |
|  | PheCA<sup>c</sup> | 965 ± 94 | 1158 ± 98 | 0.83 ± 0.12 | 3.3 ± 0.5 |
|  | LeuCA<sup>b</sup> | 42 ± 3 | 1122 ± 92 | 0.037 ±<br>0.004 | - |

<sup>a</sup>Fitted using Michaelis-Menten equation. <sup>b</sup>Fitted using linear regression. <sup>c</sup>Fitted using Hill equation.

Kinetic characterization of BSH1 (Fig. 2f-j) revealed distinct substrate preference compared to CGH. BSH1 exhibited Michaelis-Menten kinetics for GCA and TCA as substrates, with a strong preference for GCA (Fig. 2f-g, Table 1), which has been previously described.^14^ In contrast, similar studies with TyrCA and PheCA yielded sigmoidal kinetics (Fig. 2h-i) that were fit to the Hill equation^18^ and yielded Hill coefficients greater than 1 for these substrates (Table 1), which indicates positive cooperativity.^18^ Whereas positive cooperativity is reported for a BSH from *Enterococcus faecalis*,^18^ we observed cooperativity in a substrate-dependent manner for BSH1. LeuCA, however, showed linear kinetics (Fig. 2j) and displayed the lowest catalytic efficiency (Table 1). We also determined that BSH1 had a lower catalytic efficiency for TCA compared to TyrCA and PheCA (Table 1), demonstrating that MCBAs can be preferred substrates compared to host-conjugated BAs.

To understand the observed substrate preferences, we performed molecular docking analyses of these enzymes by modeling enzyme-substrate complexes for each substrate and enzyme. For CGH, we used a previously reported crystal structure of the apo enzyme with deoxycholate and taurine (PDB: 2BJF).^21^ As the structure of BSH1 is not yet available, we used ColabFold-AlphaFold2^19^ to compute a homotetrameric^16,21^ structure (Figs. 3a, S4). Then, we performed ligand docking^23^ of host-derived and MCBA substrates with these structures. Whereas host-conjugated BAs readily docked, only TyrCA and PheCA successfully produced binding poses for CGH and BSH1, respectively. For the remaining MCBA substrates, we used the docked MCBA substrates and modified the side chain scaffolds.^38^ Finally, we performed energy minimization^24^ to generate models of each enzyme-substrate complex.

**Figure 3.**
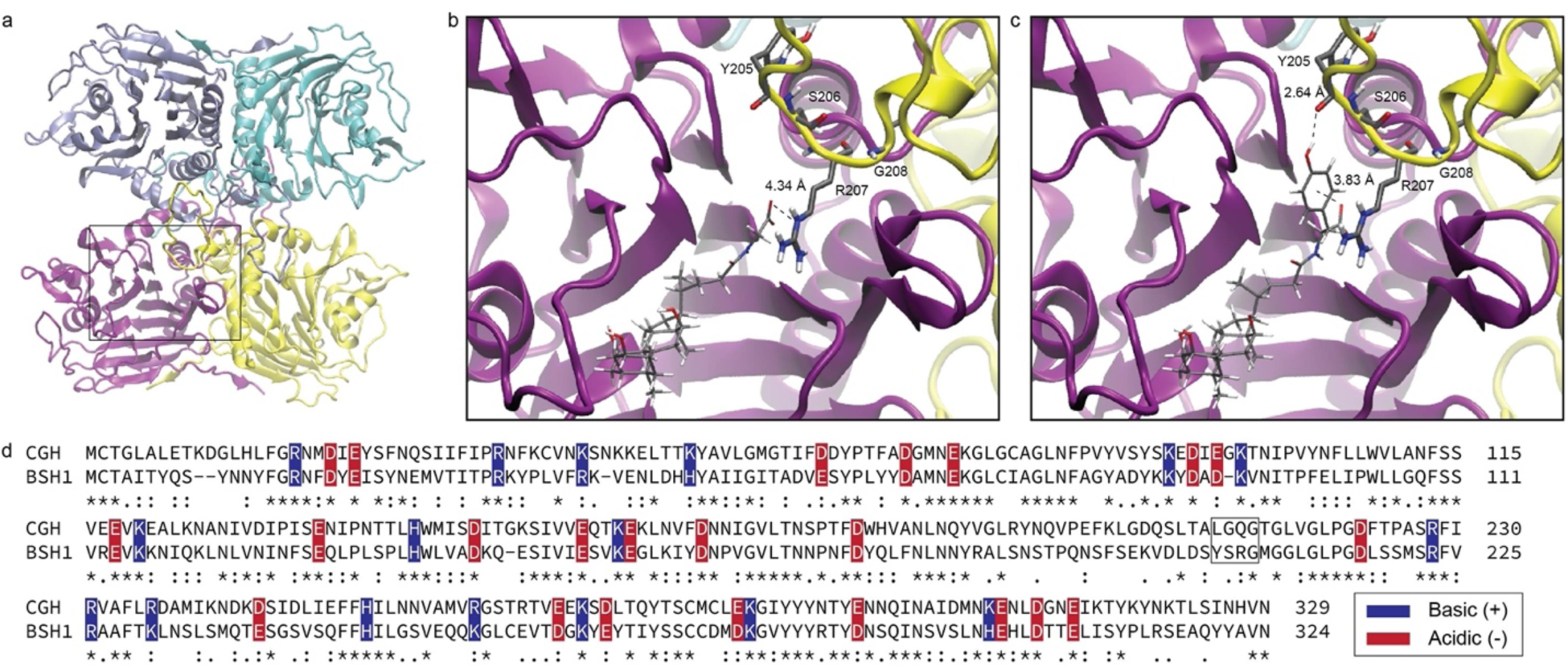
Molecular modeling analysis of *L. plantarum* BSH1. (a) Modeled homotetramer structure. Monomeric chains are colored purple, yellow, cyan, and blue. Modeled complexes for (b) GCA and (c) TyrCA, with interactions with selectivity loop (ball and stick, yellow loop) shown.

The modeled complexes of CGH place the amino acid substituent of the BA conjugate in a solvent-exposed position (Fig. S5). Whereas the polar amino acid substituent is well-solvated for host-conjugated BAs, the hydrophobic sidechain of MCBAs is exposed to solvent in these models (Fig. S5). This modeling analysis also suggests TyrCA binding is stabilized by substrate-specific H-bonding (Fig. S6), which agrees with the observed preference of CGH for TyrCA compared to PheCA (Table 1).

Atom coloring (ball and stick models): Grey (carbon), red (oxygen), blue (nitrogen), white (hydrogen). (d) Sequence alignment of *C. perfringens* CGH (CGH) and of *L. plantarum* BSH1 (BSH1). Neighboring loop residues of CGH and BSH1 are highlighted (black box). Conserved negatively- and positively-charged residues are highlighted with red and blue boxes, respectively. Conserved residues are indicated with asterisks, whereas conservatively substituted and semiconservatively substituted residues are indicated with colons and periods, respectively.

The modeled complexes of BSH1 suggested that a four amino acid loop from one monomer interacts with a bile acid substrate bound to a neighboring monomer, which includes the previously reported three amino acid selectivity loop that governs substrate preference for host-conjugated BAs (Fig. 3b-c).^16^ These models found that a conserved arginine residue, R207, is positioned over the BA conjugate amino acid (Fig. 3b-c) and is part of a loop with the sequence motif X1-S-R-X2. For GCA, the carboxylate moiety forms an electrostatic interaction with R207, which is consistent with previous structural studies (Fig. 3b).^16^ This interaction is also present for TCA (Fig. S7a) but is attenuated for LeuCA (Fig. S7b).

Interestingly, R207 formed cation-π interactions with the aryl rings of TyrCA (Fig. 3c) and PheCA (Fig. S7c) in the BSH1 models, and TyrCA engaged in an additional hydrogen bond with Y204 (Fig. 3c). To characterize these cation-π interactions, we determined geometric parameters (Fig. S8, Table S1),^26^ which were consistent with empirically identified cationic arginine-π interactions in protein structures.^26^ To determine the importance of the cationic arginine (R207)- π interaction in conferring BSH1 substrate preference to aromatic MCBAs, we generated a gain-of-function CGH Q212R mutant and a loss-of-function BSH1 R207Q mutant (Fig. S1) and tested substrate preference for GCA, TCA, and PheCA (Fig. S9). We found that the CGH Q212R mutant exhibited increased preference for PheCA and no change for GCA and TCA compared to wild-type CGH. However, the BSH1 R207Q mutant had no difference in substrate preferences for these conjugated BAs compared to wild-type BSH1, likely due to solvation effects.^16,21^ Overall, these modeling studies suggest the observed substrate preference of BSH1 for PheCA and TyrCA is due to substrate stabilization by a cation-π interaction.

Although the structure of CGH (PDB:2BJF) used for our modeling analysis was monomeric (Fig. S5),^21^ quaternary structure may be important in substrate recognition (Fig. 3a-c).^16^ Indeed, a similar loop is observed for CGH in its homotetrameric structure.^21^ Sequence alignment^31^ of CGH and BSH1 suggest that this loop in CGH bears an overall neutral charge despite the conservation of other acidic and basic residues (Fig. 3d). This sequence analysis suggests that CGH lacks stabilizing interactions with MCBA substrates, which may contribute to its lower activity toward MCBAs compared to host-conjugated substrates (Fig. 2a-e).

As our modeling studies indicated the X1-S-R-X2 motif may be important for MCBA substrate recognition, we examined the distribution of this motif in BSHs encoded by the human gut microbiome. We obtained protein sequences for the cholylglycine hydrolase subfamily (EC 3.5.1.24) from NCBI and restricted the analysis to taxa identified in the human gut. Because this motif appears at positions 205-209 in *L. plantarum* BSH1, we searched for this motif at similar positions using bioinformatics analysis. This analysis revealed that this motif is found in 2,222 (24%) of the 9,452 sequences that we examined (Table S2). The aromatic MCBA selectivity loop sequence that we identified in BSH1, YSRG, was the most abundant motif, with YSRS and FSRG comprising a much smaller fraction of sequences (Table S3).

Although BSHs are widely distributed across gut bacterial phyla,^39,40^ BSHs with this motif were found primarily in the *Bacillota* (formerly *Firmicutes*) phylum, which is a dominant phyla in the intestines of healthy individuals, and we observed distinct clades of BSHs with this motif (Figs. S10, 4a, Table S4). Phylum-level analysis also identified BSH sequences from human gut archaea (Fig. S10) for which BSH activity has been previously identified.^39^

Within *Bacillota*, we identified BSH protein sequences with genus-level taxonomic assignments and characterized the abundances of genera representing 1% or more of these sequences (Fig. 4b). This genus-level analysis found this motif was most abundant in BSH sequences from *Enterococcus*,^18^ with other genera comprising a much smaller fraction of these sequences (Fig. 4b). We also identified this motif in genera of the *Lactobacilliceae* family^16^ and the *Clostridium* genus (Fig. 4b). The additional genera that we identified are among the most abundant in the human gut microbiome (Figure 4b).^41^ Collectively, our bioinformatic analysis suggests that BSHs capable of degrading MCBAs are distributed across dominant genera of the human gut microbiome.

**Figure 4.**
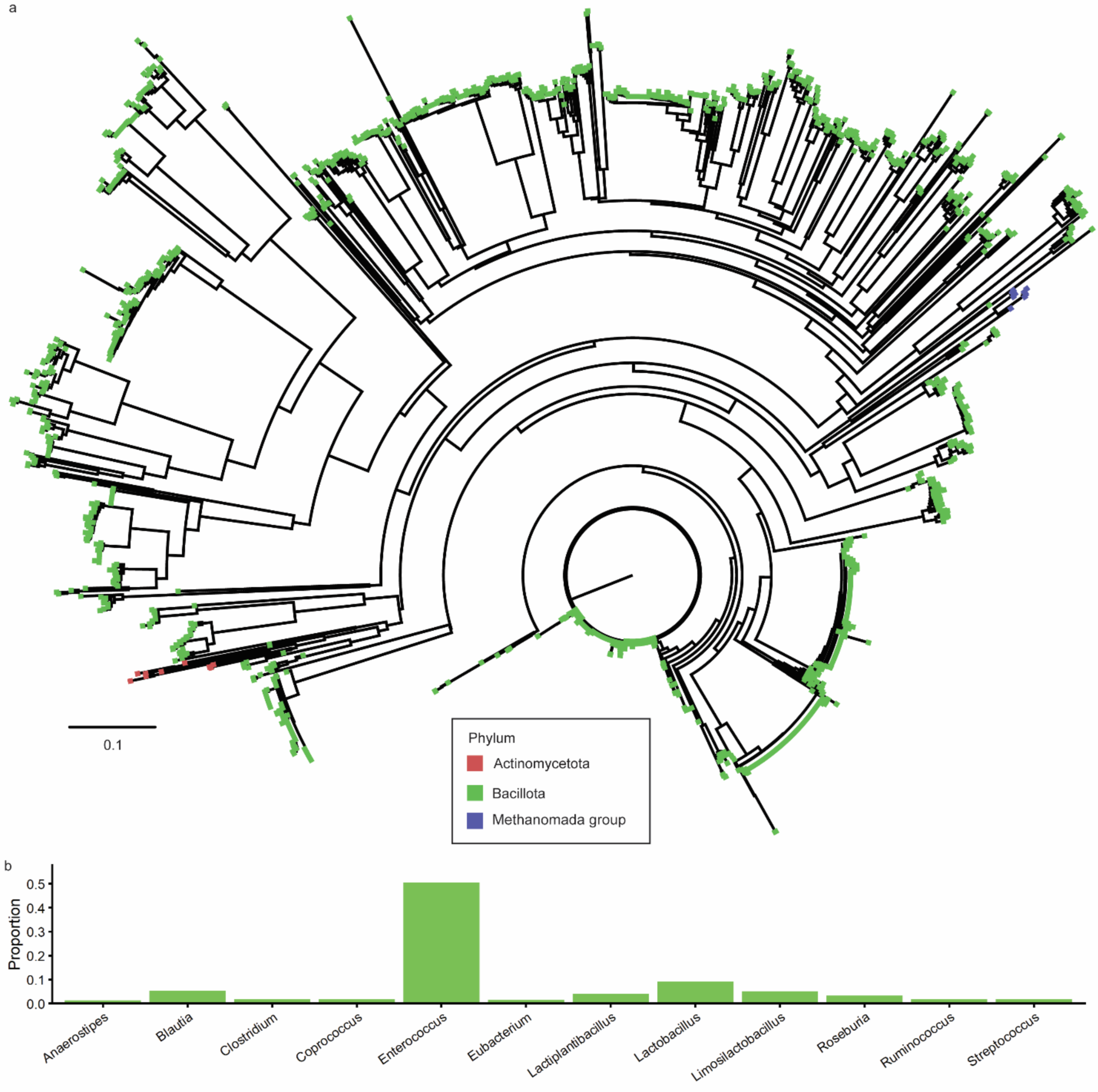
Phylogenetic analysis of BSH protein sequences from the human gut containing an aromatic MCBA selectivity loop. (a) Neighbor-joining tree of BSHs from MCBA selectivity loop, with tips colored by phylum. Tree scale represents phylogenetic distance determined by BLOSUM62^32^ score, which is based on sequence similarity where a distance of 1.0 means 100% similar. (b) Genus-level abundances for human gut microbial *Bacillota* (formerly *Firmicutes*)

BSHs containing aromatic MCBA selectivity loop. Genera accounting for 1% or more of all sequences with assigned genera are shown.

## Conclusions

In summary, BA metabolism by the gut microbiota is characterized by diverse chemical transformations and is an important process for regulating host physiology.^4^ Gut microbial BA deconjugation, catalyzed by BSHs, is critical for secondary BA metabolism in the intestines. Recent studies identified BA conjugation, which was previously thought to only occur via host-encoded enzymes,^4^ as a widespread transformation mediated by the gut microbiota to produce MCBAs.^5,6,7^ To understand the role of BSH in deconjugation of MCBAs, we biochemically characterized the substrate preference of two representative BSHs, *L. plantarum* BSH1 and *C. perfringens* CGH, for host and gut microbial cholic acid conjugates.

Our biochemical studies found that whereas host-conjugated BAs are generally preferred substrates, BSHs may have unexpected selectivity for MCBAs with specific amino acid sidechains, as was observed for *L. plantarum* BSH1. Based on molecular modeling studies, we propose that a cation-ρε interaction is responsible for the higher activity of BSH1 on TyrCA and PheCA compared to host-conjugated TCA. This molecular modeling analysis also enabled the identification of a conserved four-amino acid motif that may be responsible for this interaction, which we found is widely distributed among BSHs from dominant microbial phyla and genera in the human gut. Collectively, these results suggest that differences in BSH substrate preference may enable certain gut microbes to degrade specific MCBAs and this selective MCBA-targeted activity is widely distributed across the human gut microbiome.

## Associated content

Supporting Information is available free of charge at Protein purification, kinetics, molecular modeling, characterization of mutant BSH enzymes, phylogenetic analysis, cation-pi interaction parameters, frequency of selectivity loop motif (PDF)

## Author contributions

K.P.M. carried out the experiments and computational analyses. K.P.M. and P.V.C. conceived of the project idea, designed experiments, analyzed data, and wrote the manuscript.

## Notes

The authors declare no competing financial interests.

## Acknowledgments

This work was supported by an NIH R35 Maximizing Investigators’ Research Award for Early Stage Investigators (R35GM133501). Research in the Chang Lab is supported by a Beckman Young Investigator Award (to P.V.C.) from the Arnold and Mabel Beckman Foundation and a Sloan Research Fellowship (to P.V.C.) from the Alfred P. Sloan Foundation. K.P.M. is supported by an NIH Chemistry-Biology Interface Predoctoral Training Grant (T32GM138826). This work made use of the Cornell University NMR Facility, which is supported, in part, by the NSF through MRI award CHE-1531632. We thank the Weill Institute for Cell and Molecular Biology for additional resources.

## NCBI Accession Codes

CBAH (CGH): P54965.3 BSH1: CCC80500.1

## Author Information

### Corresponding Author

Pamela V. Chang-Department of Microbiology and Immunology, Cornell University, 930 Campus Road, Ithaca, NY 14853, United States; Department of Chemistry and Chemical Biology, Cornell University, Ithaca, NY 14853, United States; Cornell Center for Immunology, Cornell University, Ithaca, NY 14853, United States; Cornell Institute of Host-Microbe Interactions and Disease, Cornell University, 930 Campus Road, Ithaca, NY 14853, United States orcid.org/0000-0003-3819-9994; Phone: 1-607-253-4079;

### Author

Kien P. Malarney-Department of Microbiology, Cornell University, 930 Campus Road, Ithaca, NY 14853, United States; orcid.org/0000-0002-7672-4916

Complete contact information is available at:

## Notes

### Competing Interest Statement

The authors have declared no competing interest.

### Summary of Updates

I added periods to the middle initials in the author line, which were missing.

