## Supporting Information for "Electrostatic Interactions Dictate Bile Salt Hydrolase Substrate Preference"

### Contents

Figure S1. SDS-PAGE analysis of purified recombinant BSHs.

Figure S2. Linear range determination for *C. perfringens* CGH substrates.

Figure S3. Linear range determination for *L. plantarum* BSH1 substrates.

Figure S4. By-residue confidence score of predicted *L. plantarum* BSH1 structure.

Figure S5. Molecular modeling analysis of *C. perfringens* CGH.

Figure S6. Substrate-specific interactions of TyrCA and PheCA with *C. perfringens* CGH.

Figure S7. Molecular modeling analysis of *L. plantarum* BSH1.

Figure S8. Analysis of cation – π interaction between *L. plantarum* BSH1 R207 and modeled aromatic MCBAs.

Figure S9. Characterization of mutant BSH enzymes.

Figure S10. Phylogenetic analysis of BSH protein sequences from the human gut.

Table S1. Interaction parameters for cation – π interactions between *L. plantarum* BSH1 R207 and aromatic amino acid conjugates.

Table S2. NCBI protein accession numbers for BSH protein sequences examined in phylogenetic analyses (separate file).

Table S3. Frequencies of identified aromatic MCBA selectivity loop motifs within BSHs.

Table S4. Phylum-level frequencies of human gut microbial BSHs containing aromatic MCBA selectivity loop motif.

References

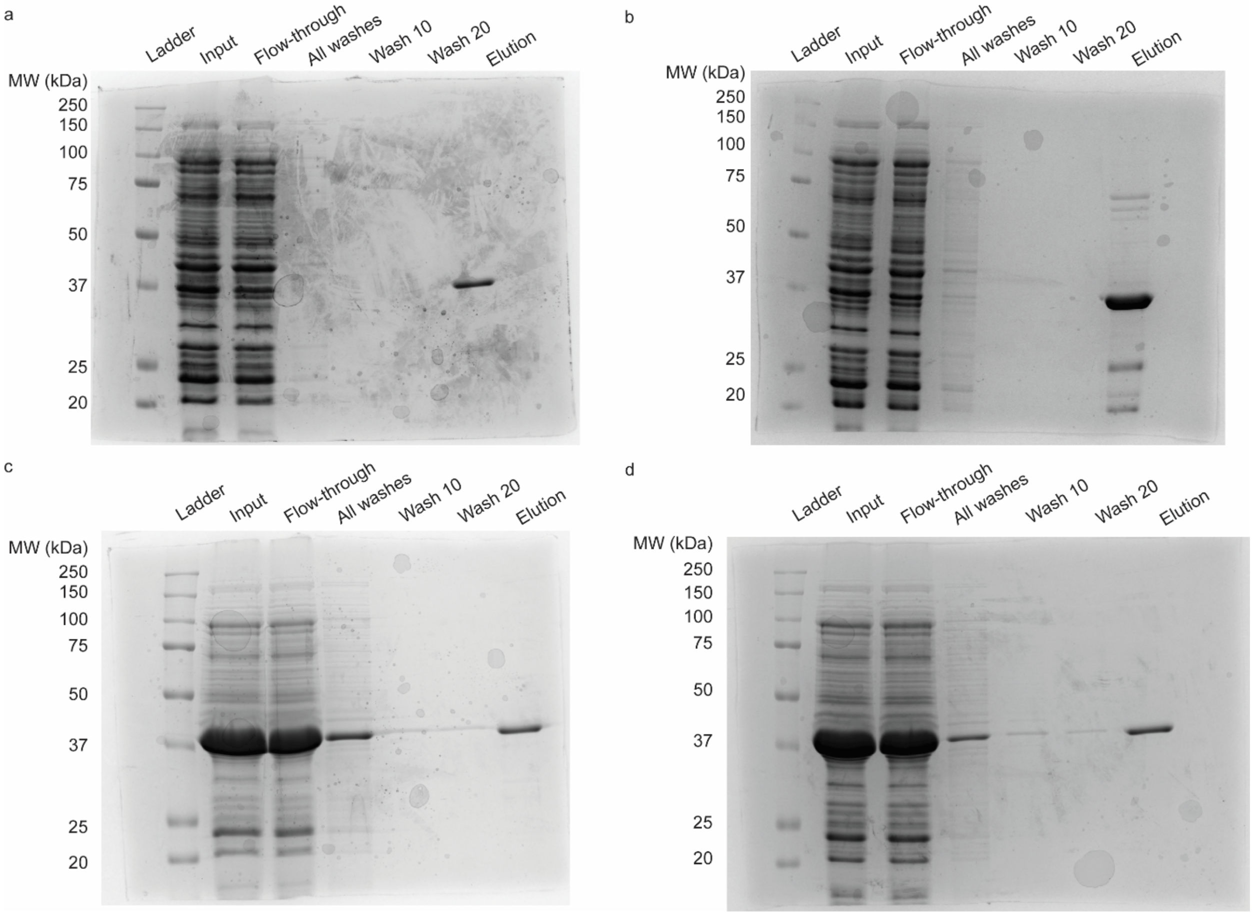


**Figure S1**. SDS-PAGE analysis of purified recombinant BSHs. Analysis of (a) wildtype *C. perfringens* CGH, (b) *C. perfringens* CGH Q212R mutant, (c) wildtype *L. plantarum* BSH1, and (d) *L. plantarum* BSH1 R207Q mutant.

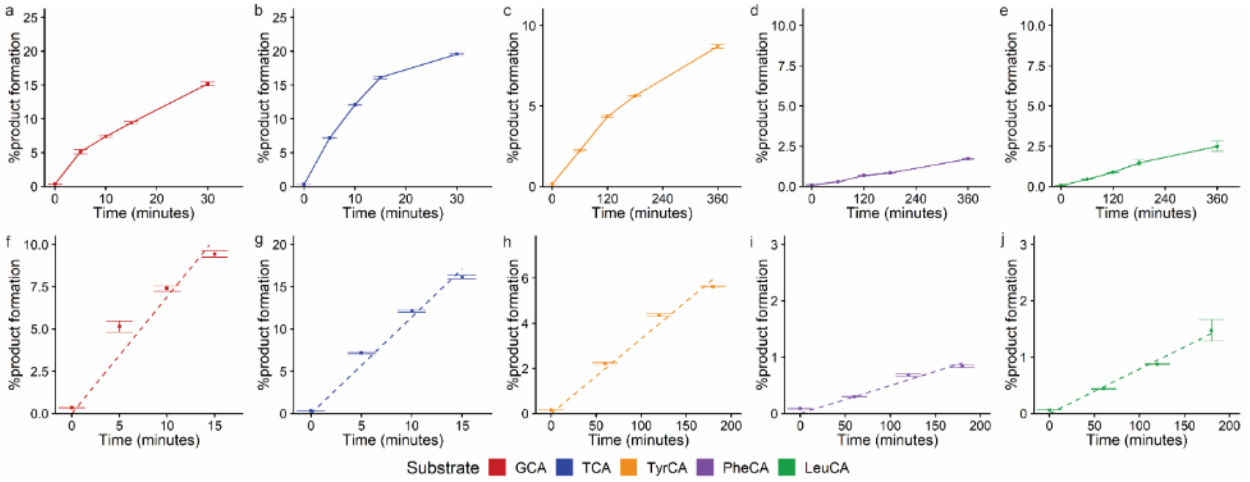


**Figure S2**. Linear range determination for *C. perfringens* CGH substrates. (a-e) Reaction progress of bile acid conjugate hydrolysis. (f-j) Linear time range of hydrolysis. BSH (a) (9.7 nM) and (b-e) (387.8 nM) was incubated with 1 mM bile acid in phosphate buffered saline, pH = 6.2, with 10 mM DTT at 37 °C. Data are representative of three independent experiments, n=3, points = mean, error bars = standard deviation.

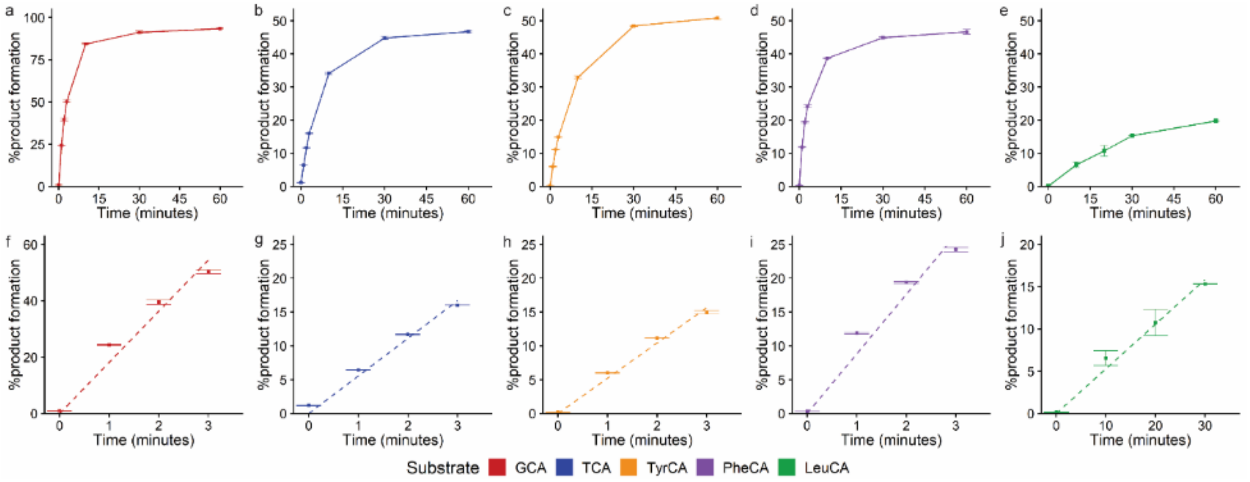


**Figure S3**. Linear range determination for *L. plantarum* BSH1 substrates. (a-e) Reaction progress of bile acid conjugate hydrolysis. (f-j) Linear time range of hydrolysis. BSH (a) (9.7 nM) and (b-e) 193.4 was incubated with 1 mM bile acid in phosphate buffered saline, pH = 6.2, with 10 mM DTT at 37 °C. Data are representative of three independent experiments, n=3, points = mean, error bars = standard deviation.

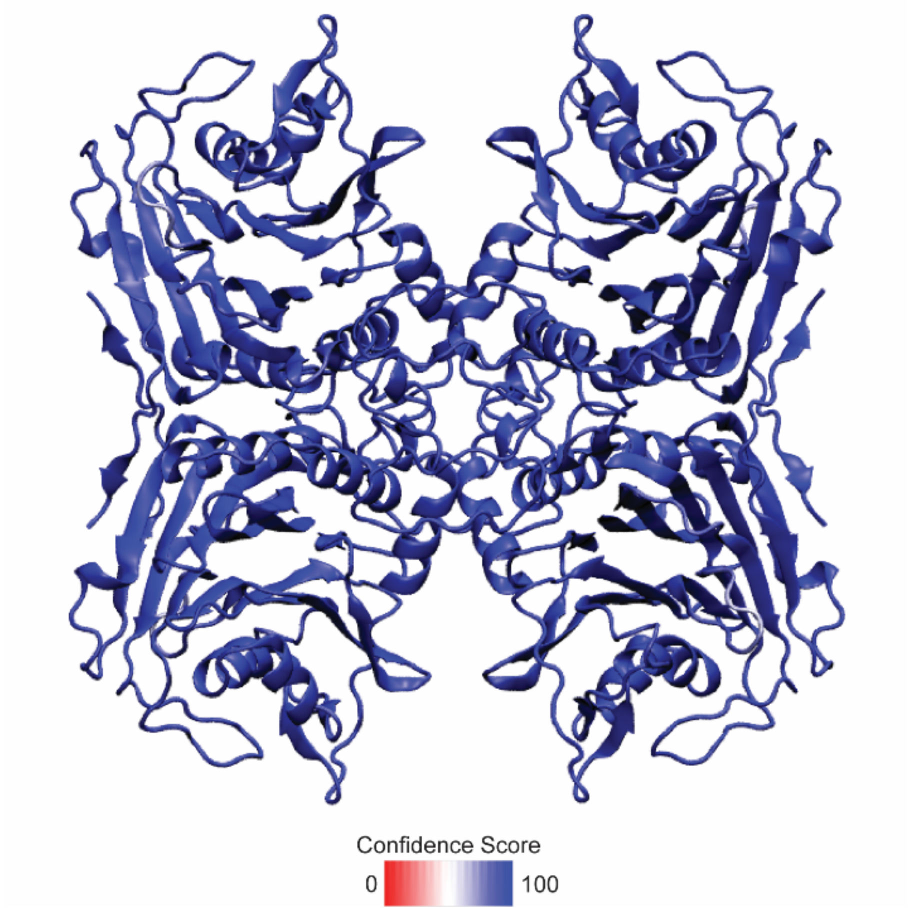


**Figure S4**. By-residue confidence score of predicted *L. plantarum* BSH1 structure using ColabFold-AlphaFold2.[3]

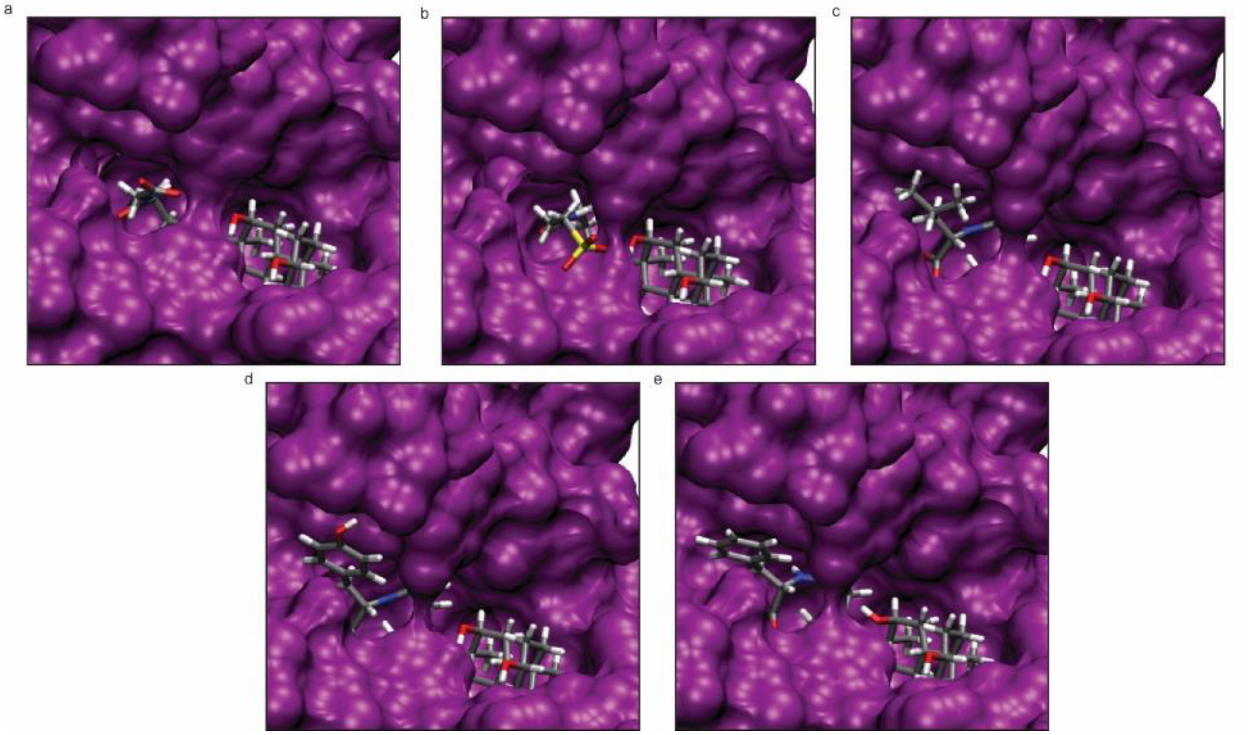


**Figure S5.** Molecular modeling analysis of *C. pefringens* CGH. Modeled complexes of *C. perfringens* CGH with (a) GCA, (b) TCA, (c) LeuCA, (d) TyrCA, and (e) PheCA. Protein is colored purple. Atom coloring (ball and stick models): Grey (carbon), red (oxygen), blue (nitrogen), yellow (sulfur), white (hydrogen).

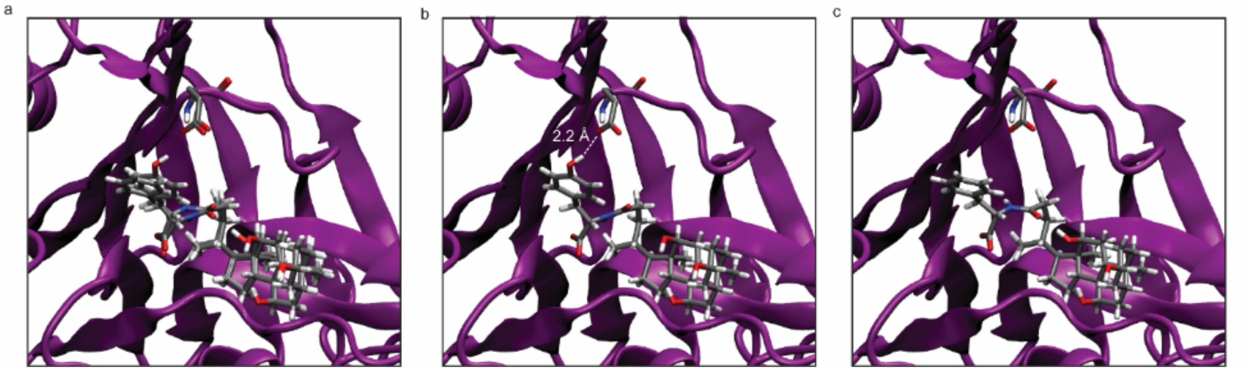


**Figure S6.** Substrate-specific interactions of TyrCA and PheCA with *C. perfringens* CGH. (a) Modeled complexes of TyrCA and PheCA, with Glu23 (ball and stick). (b) Modeled complex of TyrCA, with *para*-hydroxyl group engaged in H-bonding with Glu23 (white dashes). (c) Modeled complex of PheCA is unable to H-bond with Glu23. Protein is colored purple. Atom coloring (ball and stick models): Grey (carbon), red (oxygen), blue (nitrogen), white (hydrogen).

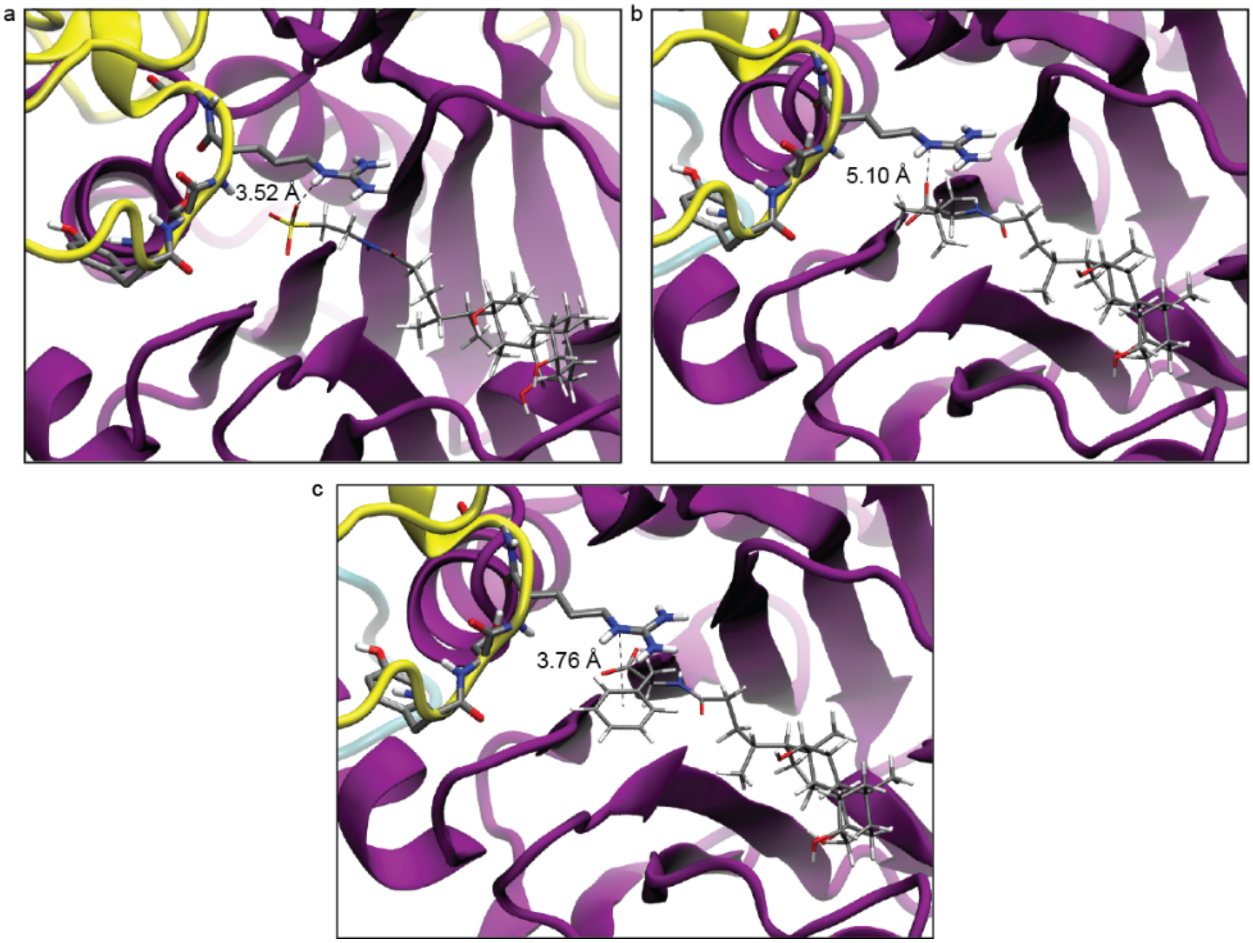


**Figure S7.** Molecular modeling analysis of *L. plantarum* BSH1. Modeled complexes of (a) TCA, (b) LeuCA, and (c) PheCA with *L. plantarum* BSH1. Electrostatic interactions with R207 are shown. Atom coloring (ball and stick models): Grey (carbon), red (oxygen), blue (nitrogen), yellow (sulfur), white (hydrogen). Relevant monomers are colored purple, yellow, and cyan.

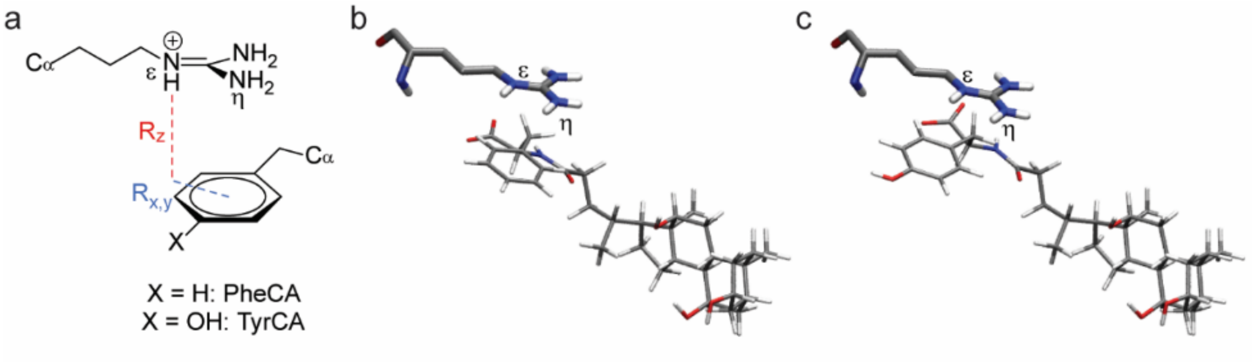


**Figure S8**. Analysis of cation – π interaction between *L. plantarum* BSH1 R207 and modeled aromatic MCBAs. (a) Definitions of distances and positions for cation – π interactions[10] between R207 and aromatic MCBAs. (b) Modeled PheCA binding and position of R207. (c) Modeled TyrCA binding and position of R207. Atom coloring (ball and stick models): Grey (carbon), red (oxygen), blue (nitrogen), white (hydrogen).

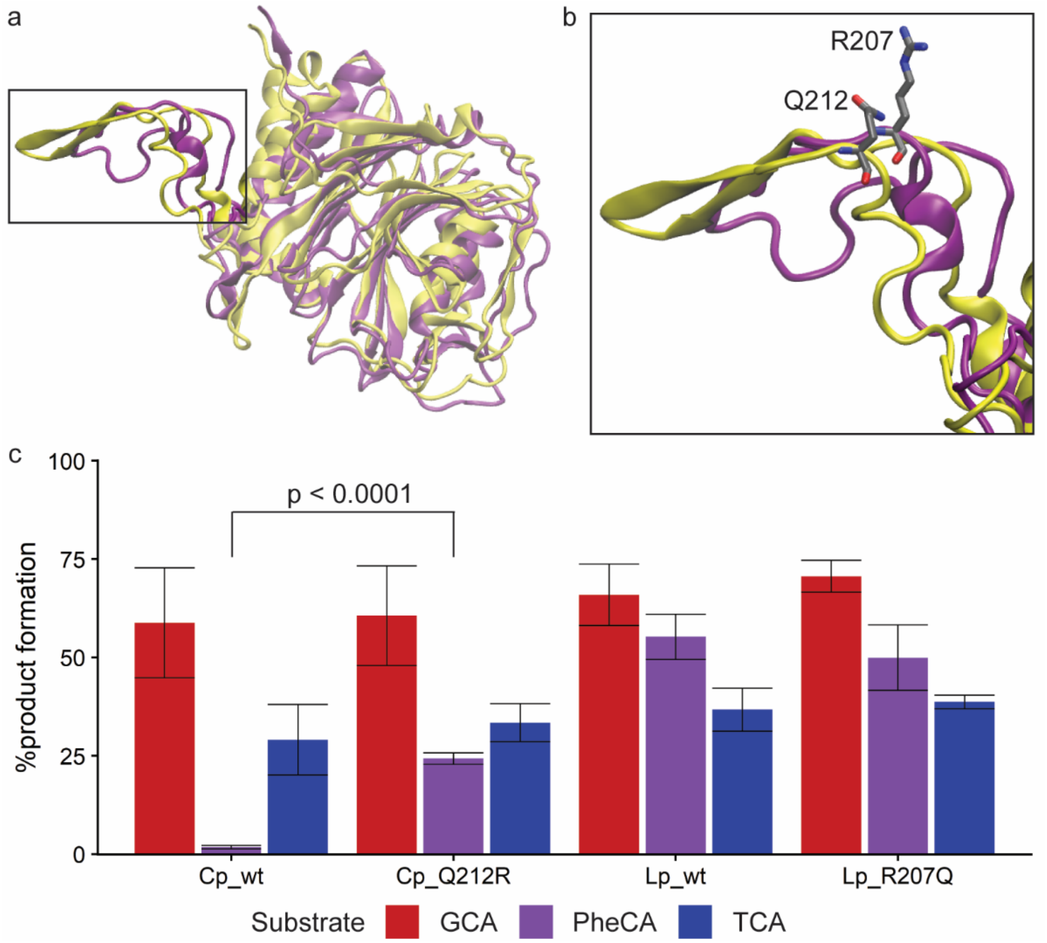


**Figure S9**. Characterization of mutant BSH enzymes. (a) Aligned monomers of *C. perfringens* CGH crystal structure (yellow) and *L. plantarum* BSH1 modeled structure (purple). (b) Selectivity loop residues R207 of *L. plantarum* BSH1 and Q212 of *C. perfringens* CGH. Atom coloring (ball and stick models): Grey (carbon), red (oxygen), blue (nitrogen). (c) Product formation after incubation of BSH (387.8 nM) with 1 mM conjugated bile acid in phosphate buffered saline, pH = 6.2, with 10 mM DTT for 5 h at 37 °C. Data are shown as mean ± standard deviation (average of $n$ = 4 independent experiments). Significant differences between wildtype and mutant BSHs were determined by Welch's t-test.

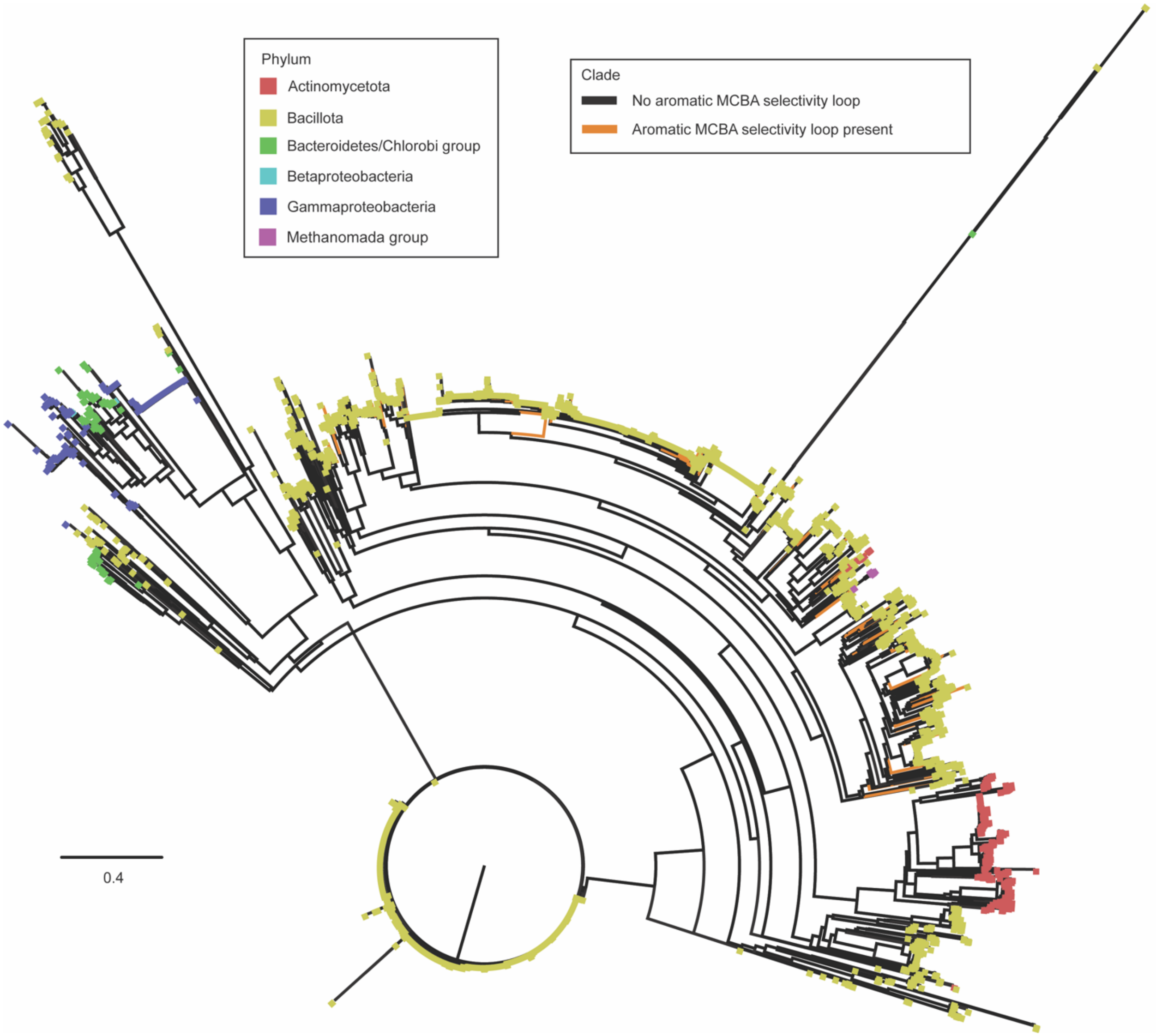


**Figure S10**. Phylogenetic analysis of BSH protein sequences from the human gut, colored by phylum (tips) and aromatic MCBA selectivity loop (clade). Tree scale represents phylogenetic distance determined by BLOSUM62[17] score, which is based on sequence similarity where a distance of 1.0 means 100% similar.

**Table S1**. Interaction parameters for cation – π interactions between *L. plantarum* BSH1 R207 and aromatic amino acid conjugates.

| Substrate | R207 nitrogen | $R_{x,y}$ (Å) | $R_z$ (Å) | Guanidinium – aryl interplane angle (°) |
|---|---|---|---|---|
| PheCA | ε | 2.26 | 3.18 | 7.39 |
| PheCA | η | 1.32 | 3.52 | |
| TyrCA | ε | 1.33 | 3.59 | 8.48 |
| TyrCA | η | 2.57 | 3.88 | |

**Table S2**. NCBI protein accession numbers for BSH protein sequences examined in phylogenetic analyses (separate file).

**Table S3**. Frequencies of identified aromatic MCBA selectivity loop motifs within BSHs.

| Sequence | Frequency |
|---|---|
| FSRG | 2/2222 (0.09%) |
| YSRG | 2186/2222 (98.38%) |
| YSRS | 34/2222 (1.53%) |

**Table S4**. Phylum-level frequencies of human gut microbial BSHs containing aromatic MCBA selectivity loop motif.

| Phylum | Frequency |
|---|---|
| Actinomycetota | 9/2222 (0.4%) |
| Bacillota | 2202/2222 (99.1%) |
| Methanomada group | 11/2222 (0.5%) |
